# The positive dimension of schizotypy is associated with a reduced attenuation and precision of self-generated touch

**DOI:** 10.1101/2022.01.22.476743

**Authors:** Evridiki Asimakidou, Xavier Job, Konstantina Kilteni

## Abstract

The brain predicts the sensory consequences of our movements and uses these predictions to attenuate the perception of self-generated sensations. Accordingly, self-generated touch feels weaker than externally generated touch of identical intensity. In schizophrenia, this somatosensory attenuation is substantially reduced, suggesting that patients with positive symptoms fail to accurately predict and process self-generated touch. Here we hypothesized that a similar impairment might exist in healthy nonclinical individuals with high positive schizotypal traits. One hundred healthy participants (53 female) scored for schizotypal traits and underwent a well-established psychophysics force discrimination task to quantify how they perceived self-generated and externally generated touch. The perceived intensity of tactile stimuli delivered to their left index finger (magnitude) and the ability to discriminate the stimuli (precision) were measured. We observed that higher positive schizotypal traits were associated with reduced somatosensory attenuation and poorer somatosensory precision of self-generated touch. These effects were specific to positive schizotypy and were not observed for the negative or disorganized dimensions of schizotypy. The results suggest that positive schizotypal traits are associated with a reduced ability to predict and process self-generated tactile stimuli. Given that the positive dimension of schizotypy represents the analogue of positive psychotic symptoms of schizophrenia, deficits in processing self-generated tactile information could indicate increased liability to schizophrenia.

## Introduction

Distinguishing between the two causes of our sensory input – the self and the environment – is fundamental for survival. First, it enables the nervous system to detect situations that can be physically harmful for the organism and to act accordingly (Brooks and Cullen, 2019; Crapse and Sommer, 2008; McNamee and Wolpert, 2019): for example, the touch of a spider crawling up one’s arm (externally generated touch) elicits a dramatically different response from the same touch applied by one’s other hand (self-generated touch). Second, this distinction is a prerequisite for maintaining our self-consciousness and consequently our mental health because it allows us to delimit our own intentions, sensations, actions, thoughts and emotions from those of others and perceive ourselves as independent human entities (Blakemore and Frith, 2003; Frith, 2005a; Leptourgos and Corlett, 2020). For example, we do not mistake our thoughts for the voices of other people we simultaneously have conversation with because we attribute the cause of our thoughts to ourselves (self-generated ‘voices’) and the cause of the voices we hear to the other people (externally generated voices).

But how do we make this distinction? One strategy of the brain is to use internal forward models to predict the sensory consequences of the movement (corollary discharge) using a copy of the motor command (efference copy) (Blakemore et al., 2000b; McNamee and Wolpert, 2019; Wolpert and Flanagan, 2001). These predictions are essential for the fast, online control of our movements because they allow the brain to estimate our body state and make corrections despite the inherent delays in the sensory system (Davidson and Wolpert, 2005; Kawato, 1999; McNamee and Wolpert, 2019; Shadmehr et al., 2008). Importantly, however, these predictions allow the brain to differentiate between self-generated and externally generated sensations: accordingly, those sensations that match the sensory predictions are self-generated, while those that deviate from the predicted ones, or have not been predicted, are attributed to external causes (Frith, 2012). Moreover, the brain uses these predictions to attenuate the intensity of the self-generated signals, thereby amplifying the difference between self-generated and externally generated information (Bäß et al., 2008; Blakemore et al., 2000b; Gentsch and Schütz-Bosbach, 2011; Kilteni et al., 2020). In the tactile domain, this attenuation manifests in perceiving self-generated touch as being weaker than externally generated touch of the same intensity (Bays et al., 2005; Bays and Wolpert, 2008; Sarah Jayne Blakemore et al., 1999; Kilteni et al., 2021, 2020, 2018; Kilteni and Ehrsson, 2020b, 2020a, 2017a, 2017b; Lalouni et al., 2020; Shergill et al., 2003) and in yielding weaker activity in the secondary somatosensory cortex and the cerebellum (Blakemore et al., 1998; Kilteni and Ehrsson, 2020a) and increased functional connectivity between the two areas (Kilteni and Ehrsson, 2020a). Somatosensory attenuation has been shown across a wide age range (18-88 years old) (Wolpe et al., 2016), and it is considered one of the reasons why we cannot tickle ourselves (Blakemore et al., 2000b; Leavens and Bard, 2016; Weiskrantz, L., Elliot, J. & Darlington, 1971).

In contrast to healthy individuals, patients with schizophrenia show an impairment in attenuating self-generated tactile sensations. Specifically, patients show significantly less attenuation of self-generated forces at the behavioral level (Shergill et al., 2005) and do not exhibit attenuation of somatosensory cortical activation for self-generated forces as healthy controls do (Shergill et al., 2014). Moreover, patients with positive symptoms, such as auditory hallucinations and delusions of control, often fail to attenuate self-generated touch and perceive such touches as if they were externally generated (Blakemore et al., 2000a). Critically, this failure of attenuation is positively correlated with the severity of their hallucinations: the more severe the hallucinations, the lower the somatosensory attenuation (Shergill et al., 2014).

These findings have supported the neuropsychiatric view that the positive symptoms of schizophrenia can be explained by a deficit in predicting and processing self-generated sensations (Frith, 2005b, 2019). Such a deficit should hinder the distinction between self-generated and externally generated sensations (Fletcher and Frith, 2009), reduce the sense of agency (Leptourgos and Corlett, 2020; Poletti et al., 2019), and produce perceptual aberrations (Frith et al., 2000), including delusions of control (Frith, 2005a) and auditory hallucinations (Poletti et al., 2019). Consequently, schizophrenia is tightly linked to an atypical perception of self-generated sensations but not externally generated sensations. Indeed, despite the heterogeneity of its symptoms, schizophrenia has been primarily described as a disorder of the sense of self (Park and Baxter, 2022; Postmes et al., 2014; Sass and Parnas, 2003), and self-disorders have been shown to constitute a crucial, trait-like phenotype of the schizophrenia spectrum (Henriksen et al., 2021).

If the positive symptoms of schizophrenia are intrinsically linked to deficits in predicting and processing self-generated somatosensation, then a similar relationship should exist between positive schizotypy and impaired prediction and processing of self-generated somatosensation in nonclinical individuals. Importantly, this approach circumvents many of the methodological confounds arising from patient studies, such as antipsychotic treatment, hospitalization, and disease chronicity, that the patient groups are typically subjected to (Fervaha and Remington, 2013). Schizotypy, or psychosis-proneness, describes subclinical psychosis-like symptoms or personality characteristics, including peculiar beliefs, unusual sensory experiences and odd behavior (Meehl and Prologue, 1990; Raine, 1991), that apply to the general population (Barrantes-Vidal et al., 2015; Kwapil and Barrantes-Vidal, 2015; Nelson et al., 2013; Racioppi et al., 2015; Thomas et al., 2019; Van Os et al., 2000). Schizotypal traits are presumed to originate from the same combination of genetic, neurodevelopmental and psychosocial factors as schizophrenia (Andreasen, 1999; Debbané et al., 2015; Ettinger et al., 2014; Keshavan MS., 1997; Lenzenweger, 2006; Meehl PE., 1962; Miller P, Byrne M, Hodges A, Lawrie SM, Owens DG, 2002; Rado, 1956; Weinberger, 1987), they lie on a continuum with schizophrenia (Nelson et al., 2013) and are considered a valid phenotypic indicator for the liability to psychosis spectrum disorders and for understanding the underlying psychopathology (Barrantes-Vidal et al., 2015; Ettinger et al., 2014; Kwapil and Barrantes-Vidal, 2015; Thomas et al., 2019). Similar to schizophrenia symptom clusters, schizotypy consists of three dimensions, positive, negative and disorganized (E. Fonseca-Pedrero et al., 2018; Nelson et al., 2013; Thomas et al., 2019), that broadly correspond to the positive (*e.g*., hallucinations and delusions), negative (*e.g*., alogia and apathy) and disorganized symptoms of schizophrenia (*e.g*., thought disorder and bizarre behavior) (Kwapil and Barrantes-Vidal, 2015; Liddle, 1987; Raine et al., 1994; Reynolds et al., 2000; Rossi and Daneluzzo, 2002; Stuart et al., 1999; Wuthrich and Bates, 2006).

Here, we investigated the relationship between schizotypal traits and the perception of self-generated and externally generated somatosensation in 100 healthy individuals. We hypothesized that high positive schizotypy would be associated with reduced somatosensory attenuation and lower precision of self-generated touch.

## Methods and Materials

### Participants

The data of one hundred and two participants were used in the present study. Current or history of psychological or neurological conditions, as well as the use of any psychoactive drugs or medication, were criteria for exclusion. All participants reported being completely healthy without neurological or psychiatric disorders or taking any medication to treat such conditions. Our sample size was based on two previous studies that assessed the relationship between schizotypy and tactile perception in samples consisting of non-clinical individuals (Lenzenweger, 2000; Whitford et al., 2017). The data were pooled from three studies, all including the same psychophysics task and schizotypy measure, and identical experimental conditions. Two participants were excluded because of missing data in the schizotypy measure. Thus, the final sample consisted of one hundred (100) adults (53 women and 47 men; 91 right-handed, 5 ambidextrous and 4 left-handed; age range: 18-40 years). Handedness was assessed using the Edinburgh Handedness Inventory (Oldfield, 1971). All participants provided written informed consent, and the Swedish Ethical Review Authority (https://etikprovningsmyndigheten.se/) approved all three studies (#2020-03647, #2020-03186, #2020-05457).

### Psychophysical task

The psychophysical paradigm was a two-alternative forced choice force-discrimination task that has been extensively used to assess somatosensory attenuation (Bays et al., 2006, 2005; Kilteni et al., 2021, 2020, 2019; Kilteni and Ehrsson, 2020b). Participants sat comfortably on a chair and rested their left hand, palm up, with the left index finger placed inside a molded support (**Figure 1**). Their right hand and forearm were placed on top of boxes, next to their left hand. In each trial, a DC electric motor (Maxon EC Motor EC 90 flat; manufactured in Switzerland) delivered two brief (100 ms) forces on the pulp of participants’ left index finger through a cylindrical probe (25 mm height) with a flat aluminum surface (20 mm diameter) attached to the motor’s lever. We refer to the first force as the *test* tap and to the second force as the *comparison* tap. The intensity of the *test* tap was set to 2 N, while the intensity of the *comparison* tap was systematically varied among seven different force levels (1, 1.5, 1.75, 2, 2.25, 2.5 or 3 N). In each trial, participants verbally indicated which tap felt stronger: the *test* tap or the *comparison* tap. A force sensor (FSG15N1A, Honeywell Inc.; diameter, 5 mm; minimum resolution, 0.01 N; response time, 1 ms; measurement range, 0–15 N) was placed within the probe to record the forces exerted on the left index finger. Moreover, a force of 0.1 N was constantly applied to the participants’ left index finger to ensure accurate force intensities.

**Figure 1.**
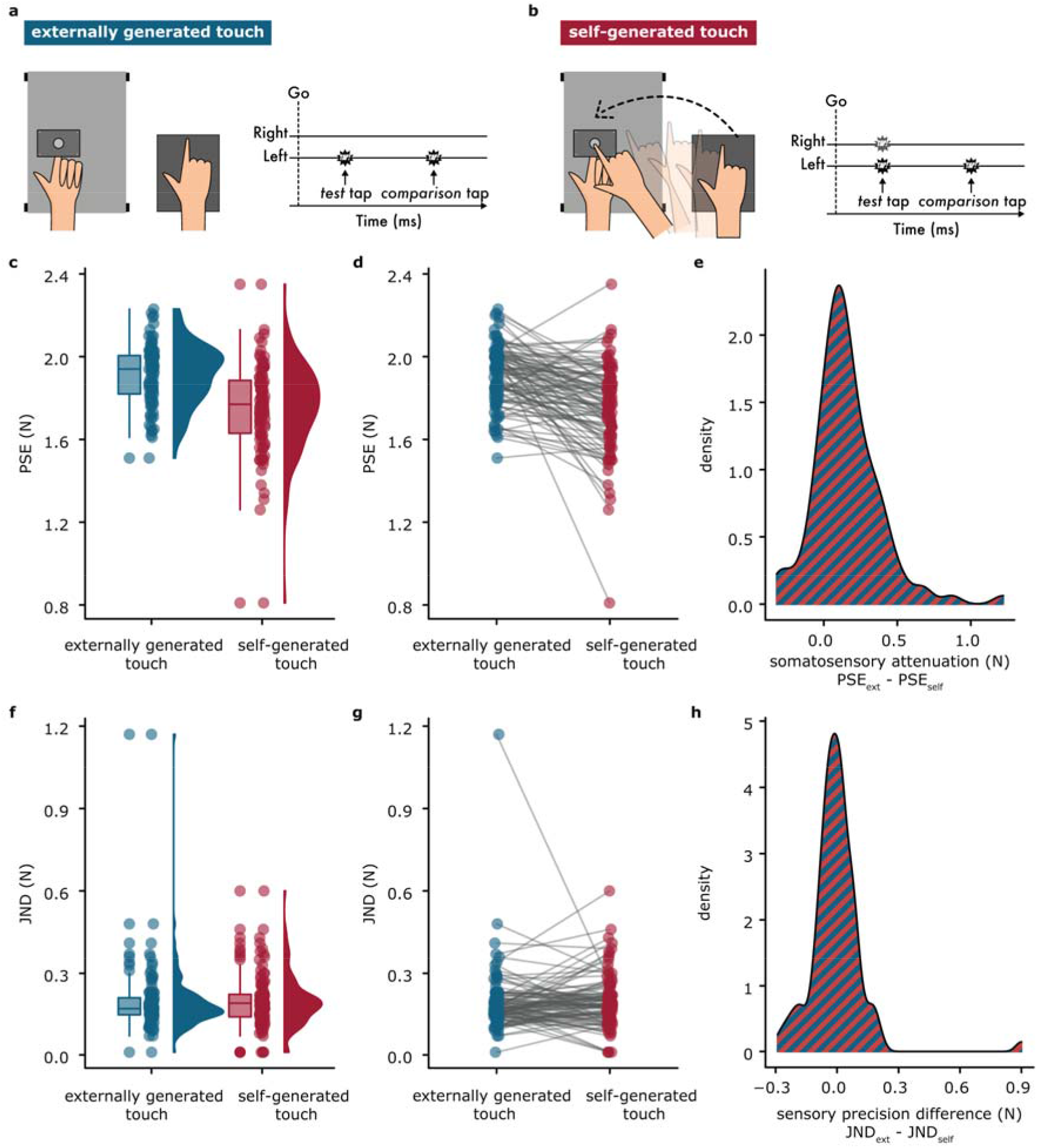
Experimental Methods and results. (**a-b**) In both conditions, the participants received two taps (*test* and *comparison* taps) on the pulp of their left index fingers, and they had to verbally indicate which felt stronger: the first or the second tap. In the externally generated touch condition **(a)**, the participants kept both of their hands relaxed while receiving the *test* tap and *comparison* taps on their left index finger. In the *self-generated touch* condition **(b)**, the participants actively tapped a force sensor with their right index finger and triggered the *test* tap on their left index finger. Then, they remained relaxed while receiving the *comparison* tap. **(c)** The boxplots show the median and interquartile ranges for the PSEs, the jittered points denote the raw data, and the violin plots display the full distribution of the data in each condition. A lower PSE value indicates a lower perceived magnitude. (**d**) Line plots illustrate the decreases in PSEs when experiencing self-generated touches compared to externally generated touches. The PSEs were significantly decreased in the *self-generated touch* condition compared to the *externally generated touch* condition. (**e**) Density plot for somatosensory attenuation (difference in the PSEs between the two conditions). Somatosensory attenuation occurred in eighty participants (80%). **(f)** The boxplots show the median and interquartile ranges for the JNDs, the jittered points denote the raw data, and the violin plots display the full distribution of the data in each condition. A lower JND value indicates a higher somatosensory precision. (**g**) Line plots illustrate the changes in JNDs when experiencing self-generated touches compared to externally generated touches. The JNDs did not significantly differ between the *self-generated touch* and *externally generated touch* conditions. (**h**) Density plot for the difference in the sensory precision between the two conditions. Forty-four participants (44%) had higher JNDs in the *externally generated touch* condition than in the *self-generated touch* condition, fifty-two participants (52%) showed the opposite pattern, and four (4%) did not change.

There were two experimental conditions, the order of which was counterbalanced across participants. In the *externally generated touch* condition, participants kept their right arm relaxed and passively received the two taps on their left index finger (**Figure 1a**). The *test* tap was delivered 800 ms after an auditory ‘go’ cue, and the *comparison* tap was delivered after a random delay (800 ms – 1500 ms) from the end of the *test* tap. In the *self-generated touch* condition, participants actively tapped with their right index finger a force sensor placed above (but not in contact with) the probe after the auditory ‘go’ cue (**Figure 1b**). They were instructed to tap the sensor neither too hard nor too softly but as strongly as when they tapped the surface of their smartphone. The tap of their right index finger triggered the *test* tap on their left index finger with an intrinsic delay of 36 ms.

Each condition consisted of 70 trials, resulting in 140 trials per participant. The order of the intensities was randomized across participants. In both conditions, the view of the pulp of the left index finger was occluded, and participants were asked to fixate on a cross placed on a wall 2 meters in front of them. Any sounds created by the motor or the participants’ taps were suppressed by administering white noise through a pair of headphones. Before the experiment, the participants were instructed to avoid balancing their responses. If the intensity of the two taps felt very similar, they were explicitly told that they had to guess. No feedback was ever provided about their performance.

### Psychophysical fits

In each condition, the participants’ responses were fitted with a generalized linear model using a *logit* link function (Equation 1)

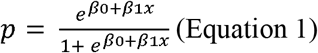

Two parameters of interest were extracted. The point of subjective equality 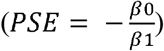 represents the intensity at which the *test* tap felt as strong as the *comparison* tap (*p* = 0.5) and quantifies the participants’ perceived intensity of the *test* tap. Subsequently, somatosensory attenuation is calculated as the difference between the PSEs of the two conditions (PSE_*external*_ – PSE_*self*_) (Bays et al., 2006, 2005; Kilteni et al., 2021, 2020, 2019; Kilteni and Ehrsson, 2020b). The just noticeable difference parameter 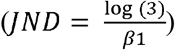 reflects the participants’ sensitivity in the psychophysics task and quantifies their somatosensory precision in each condition. The PSE and JND are independent qualities of sensory judgments (Mapp et al., n.d.).

Before fitting the responses, the values of the applied *comparison* taps were binned to the closest value with respect to their theoretical values (1, 1.5, 1.75, 2, 2.25, 2.5 or 3 N). After data collection, 951 trials out of 14000 (6.8%) were rejected, either because the intensity of the *test* tap (2 N) was not applied accurately (*test* tap < 1.85 N or *test* tap > 2.15 N) or due to missing responses.

### Schizotypal traits

After the psychophysical task, all participants completed the Schizotypal Personality Questionnaire (SPQ) (Raine, 1991). The SPQ is a 74-item self-report schizotypy assessment instrument with excellent internal reliability (*Cronbach’s alpha* = 0.91) and test-retest reliability (0.82)(Raine, 1991). It was developed on the basis of the nine features of schizotypal personality disorder, as defined by the DSM-III-R criteria (American Psychiatric Association, 1987) (Raine, 1991).

We used the three-factor model to partition the dimensions of the construct of schizotypy (E. Fonseca-Pedrero et al., 2018; Fonseca-Pedrero et al., 2014a, 2014b; Rabella et al., 2018; Raine et al., 1994; Reynolds et al., 2000; Tsaousis et al., 2015), and we calculated the total score for the cognitive-perceptual, interpersonal and disorganized factors that reflect the positive, negative and disorganized dimensions of schizotypy, respectively. There has been discussion as to whether schizotypy constitutes a continuous or a categorical construct (Kwapil and Barrantes-Vidal, 2015; Lemaitre et al., 2016; Lenzenweger, 2015; Mason, 2014; Whitford et al., 2017). In line with the predominant conceptualization of schizotypy as a continuous variable within the general population (Claridge, 1994; Kwapil and Barrantes-Vidal, 2015; Nelson et al., 2013; Van Os et al., 2000), our main analysis comprised treating positive schizotypal traits as a continuous variable across the entire sample. Nonetheless, to attain methodological rigor and to account for both notions, we performed a secondary analysis treating schizotypy as a categorical variable. Given the absence of established cut-off values for the SPQ estimates, we split the sample into 3 subgroups based on their positive schizotypal traits (*low*, *medium*, *high*). This approach was deemed appropriate to discern the differences between the two extremes (i.e., *low* and *high*).

### Statistical Analysis

Data were analyzed using R (Team, 2019) and JASP (JASP and JASP Team, 2019). Data normality was assessed using the Shapiro–Wilk test, and planned comparisons were subsequently made using parametric (independent or paired t-test) or nonparametric (Mann–Whitney or Wilcoxon) statistical tests. For each test, 95% confidence intervals (*CI^95^*) are reported. Depending on the data normality, effect sizes are given by Cohen’s *d* or by the matched rank biserial correlation *rrb*. Correlations were tested using Spearman correlation coefficients given that the data were not normally distributed. A Bayesian factor analysis was carried out for all statistical comparisons (default Cauchy priors with a scale of 0.707) and correlations (Kendall’s tau-b) to provide information about the level of support for the null hypothesis compared to the alternative hypothesis (*BF_01_*) given the data. All statistical tests were two-tailed.

## Results

### Somatosensory attenuation and precision across the entire sample

The PSE was significantly lower in the *self-generated touch* condition than in the *externally generated touch* condition across the entire sample: *n* = 100, *V* = 625, *p* < 0.001, *CI^95^* = [−0.185, −0.105], *rrb* = −0.747, *BF_01_* < 0.001 (**Figure 1c, d**). This indicates that self-generated touches felt weaker than externally generated touches of identical intensity, replicating previous findings (Bays et al., 2006, 2005; Kilteni et al., 2021, 2020, 2019; Kilteni and Ehrsson, 2020b). One participant had an extreme PSE value in the *self-generated touch* condition, as shown in **Figure 1c**. When removing the value, the same results were obtained: *n* = 99, *V* = 625, *p* < 0.001, *CI^95^* = [−0.180, −0.105], *rrb* = −0.742, *BF_01_* < 0.001. Attenuation was observed in 80% of participants in the cohort (**Figure 1e**).

In contrast to the PSEs, the JNDs did not significantly differ between the two conditions: *n* = 100, *V* = 2592, *p* = 0.335, *CI^95^* = [−0.01, 0.03], *rrb* = 0.113 (**Figure 1f, g**). This was strongly supported by a Bayesian analysis (*BF_01_* = 5.417) and indicates that self-generated and externally generated touches were perceived with similar sensory precision, in line with previous studies (Kilteni et al., 2021; Kilteni and Ehrsson, 2020b). One participant had an extreme JND value in the *externally generated touch* condition, as shown in **Figure 1f.** When removing the value, the same results were obtained: *n* = 99, *V* = 2592, *p* = 0.247, *CI^95^* = [−0.005, 0.03], *rrb* = 0.137, *BF_01_* = 3.480. As seen in **Figure 1h**, approximately half of the participants increased and half decreased their JNDs between the conditions (44% increased, 52% decreased, 4% remained unchanged).

No significant correlation was observed between the PSEs and JNDs in either the *self-generated touch* condition (*n* = 100, *rho* = 0.079, *p* = 0.437) or in the *externally generated touch* condition (*n* = 100, *rho* = 0.046, *p* = 0.647), and this was strongly confirmed by a Bayesian analysis (*BF_01_* = 5.452 for the *self-generated touch* condition, and *BF_01_* = 6.560 for the *externally generated touch* condition). This corroborates that sensory magnitude (PSE) and precision (JND) are independent measures (Mapp et al., n.d.), and is in line with previous findings (Kilteni and Ehrsson, 2020b). All individual fits are shown in **Figure S1**.

### Schizotypal traits and somatosensory attenuation

**Figure 2a-d** shows the distribution of the total SPQ scores (*μ* = 20.87, *σ* = 12.165, range = 0-53, *Cronbach’s alpha* = 0.821), as well as those of the cognitive-perceptual, interpersonal and disorganized factors in our sample (**Supplemental Table S1, Supplemental Figure S2, Supplemental Text S1**). Our schizotypy distributions were very similar to those of previous studies using random sampling methods, both in terms of mean and variability (*e.g*., (Eduardo Fonseca-Pedrero et al., 2018; Whitford et al., 2017)). Confirming our first hypothesis, we observed a negative correlation between somatosensory attenuation and schizotypal traits (*n* = 100, *rho* = −0.215, *p* = 0.031, *BF_01_* = 0.865) (**Figure 2e**), which was driven by the scores of the cognitive-perceptual factor (*i.e*., the positive dimension of schizotypy) (**Figure 2f**): *n* = 100, *rho* = −0.259, *p* = 0.009, *BF_01_* = 0.243. This means that the higher the positive schizotypal traits of the participants, the lower their somatosensory attenuation. The individual PSEs did not significantly correlate with the positive schizotypy (*self-generated touch* condition: *n* = 100, *rho* = −0.097, *p* = 0.335; *externally generated touch* condition: *n* = 100, *rho* = −0.180, *p* = 0.074). The absence of these significant correlations was supported by a Bayesian analysis (*BF_01_* = 4.794 for the *self-generated touch* condition and *BF_01_* = 1.502 for the *externally generated touch* condition), indicating that positive schizotypal traits are associated with the perceived *difference* between the intensities of a self-generated and an externally generated touch (*i.e*., somatosensory attenuation). Critically, the relationship between attenuation and schizotypy was found only for the positive schizotypy and not for the negative (*i.e*., interpersonal factor) (*n* = 100, *rho* = −0.179, *p* = 0.074) (**Figure 2g**) or the disorganized dimension (*i.e*., disorganized factor) (*n* = 100, *rho* = −0.106, *p* = 0.294) (**Figure 2h**), and a Bayesian analysis further supported the absence of these relationships (*BF_01_* = 1.552 for the negative and *BF_01_* = 4.337 for the disorganized dimension).

**Figure 2.**
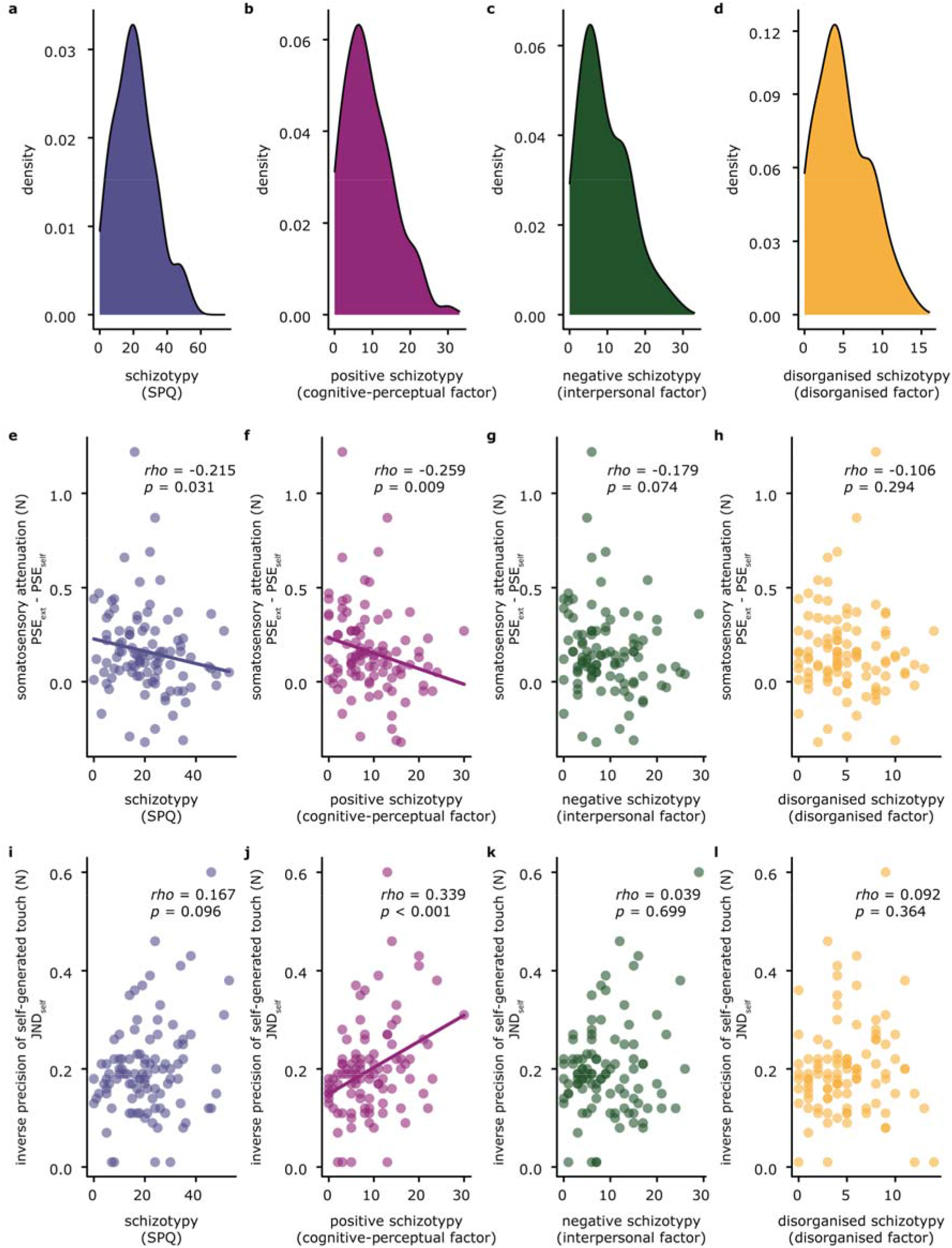
Schizotypal traits and somatosensory attenuation and precision. **(a-d)** Density plots of the Schizotypal Personality Questionnaire (SPQ) scores (possible score ranges: total, 0-74; cognitive-perceptual, 0-33; interpersonal, 0-33; disorganized, 0-16). The sample had comparable levels of positive, negative and disorganized schizotypy (**Supplemental Text S1**) and the scores covered almost the entire range of possible responses (**Supplemental Table S1**). **(e-h)** Correlations between the Schizotypal Personality Questionnaire (SPQ) scores and somatosensory attenuation. The correlations were significant only between somatosensory attenuation and the SPQ full scores, as well as between somatosensory attenuation and the cognitive-perceptual factor (positive dimension of schizotypy). Regression lines are shown for illustrative purposes only. **(i-l)** Correlations between the Schizotypal Personality Questionnaire (SPQ) scores and the inverse somatosensory precision of self-generated touch (JND). Note that the y-axis displays the JNDs (*i.e*., the inverse somatosensory precision). The correlations were significant only between somatosensory precision and the cognitive-perceptual factor (positive dimension of schizotypy). Regression lines are shown for illustrative purposes only.

### Schizotypal traits and somatosensory precision

Confirming our second hypothesis, we observed a positive correlation between the JND of self-generated touch and positive schizotypal traits (*n* = 100, *rho* = 0.339, *p* < 0.001, *BF_01_* = 0.018) (**Figure 2j**), which effectively is a negative correlation between the somatosensory precision of self-generated touch and positive schizotypal traits. In other words, the higher the positive schizotypal traits of the participants, the lower their somatosensory precision of self-generated touch. In contrast, the somatosensory precision for externally generated touch did not correlate with positive schizotypy (*n* = 100, *rho* = 0.114, *p* = 0.257, *BF_01_* = 3.639), suggesting that positive schizotypy does not generically influence the precision with which touch is perceived but only that of self-generated touch. Finally, the somatosensory precision of self-generated touch significantly correlated only with positive schizotypy but not with the full SPQ (*n* = 100, *rho* = 0.167, *p* = 0.096) (**Figure 2i**), or the negative (*n* = 100, *rho* = 0.039, *p* = 0.699) (**Figure 2k**) or disorganized dimension (*n* = 100, *rho* = 0.092, *p* = 0.364) (**Figure 2l**). As above, the Bayesian analysis strongly supported the absence of these relationships (*BF_01_* = 6.600 for the negative dimension and *BF_01_* = 4.682 for the disorganized dimension).

### Schizotypy as a categorical variable

Finally, we treated positive schizotypal traits as a categorical variable by dividing our sample into 3 subgroups with equal number of participants: the low (*n_low_ = 34*), medium (*n_med_* = 33) and high (*n_high_* = 33) positive schizotypy groups (**Figure 3a**).

**Figure 3.**
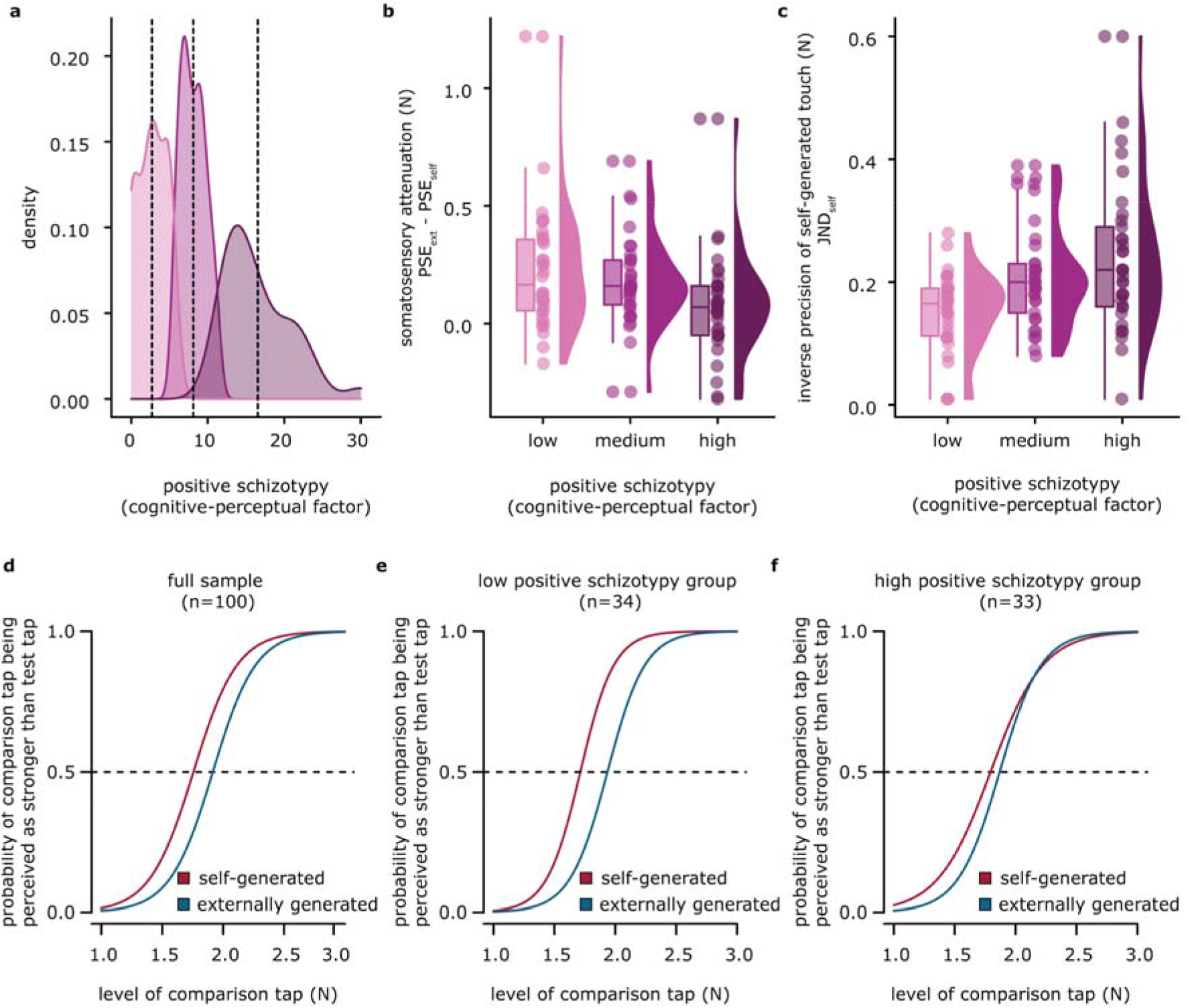
Somatosensory attenuation and precision in individuals with low, medium, and high positive schizotypal traits. **(a)** Density plots for the three schizotypy subgroups of our sample. Vertical dotted lines indicate the mean of each subgroup. **(b)** The boxplots show the median and interquartile ranges for somatosensory attenuation, the jittered points denote the raw data, and the violin plots display the full distribution of the data in each group. The high schizotypy group showed significantly less somatosensory attenuation than the low schizotypy group. **(c)** The boxplots show the median and interquartile ranges for the JND in the *self-generated touch* condition, the jittered points denote the raw data, and the violin plots display the full distribution of the data in each group. The high schizotypy group showed significantly less somatosensory precision (significantly higher JND) than the low schizotypy group. **(d)** Group psychometric fits using the total sample. The fits for each condition were generated using the mean PSE and the mean JND across participants. The leftward shift of the curve for the *self-generated touch* condition with respect to the curve for the *externally generated touch* condition illustrates that self-generated touch is perceived as weaker than externally generated touch. No change is visible in the slopes of the curves, as their JNDs did not significantly differ. **(e-f)** Group psychometric fits for the low (**e**) and the high (**f**) schizotypy groups. The high schizotypy group shows a substantially smaller shift in the curves between the *self-generated* and *externally generated* touch conditions (*i.e*., less attenuation) and a flatter slope for the *self-generated touch* condition (*i.e*., higher JND).

Somatosensory attenuation decreased from low to high schizotypy (**Figure 3b)**, yielding a significant difference between the two extremes (*n_low_ = 34, n_high_* = 33, *W* = 770, *p* = 0.009, *CI^95^* = [0.030, 0.230], *rrb* = 0.373, *BF_01_* = 0.280). In contrast, the JND increased from low to high schizotypy (*n_low_ = 34, n_high_* = 33, *t(48.7)* = −3.626, *p* < 0.001, *CI^95^* = [−0.133, −0.038], Cohen’s *d* = −0.89, *BF_01_* = 0.018) (**Figure 3c**, **Supplemental Figures S3**-**S4**, **Supplemental Text S2**).

**Figure 3d-f** illustrates these effects for the entire sample (**Figure 3d)**, the low (**Figure 3e)** and the high positive schizotypy subgroups (**Figure 3f)**. In the entire sample, the curve shifted to the left for the *self-generated touch* condition compared to the *externally generated touch* condition without any changes in the slope; thus, self-generated touch felt weaker than external touch, but they were perceived with similar precision (**Figure 3d**). Critically, as seen in **Figures 3e** and **3f**, the high positive schizotypy group showed less of a shift between the PSEs in the *self-generated* and *externally generated touch* conditions (less attenuation) and a flatter curve in the *self-generated touch* condition (higher JND) compared to the low schizotypy group.

## Discussion

The present study has two main findings. First, individuals with higher positive schizotypal traits exhibited less attenuation of their self-generated touch than individuals with low positive schizotypal traits. This result strongly mirrors previous clinical findings of reduced somatosensory attenuation in patients with schizophrenia (Blakemore et al., 2000a; Shergill et al., 2014, 2005). This is also in line with earlier observations that nonclinical individuals with high schizotypy subjectively rate self-generated touch as more ticklish (Lemaitre et al., 2016) and intense (Whitford et al., 2017) than those with low schizotypy. Second, our experimental task (i.e., the force-discrimination task) enabled the measurement, not only of the perceived magnitude but also of that of somatosensory precision and consequently, the assessment of its relationship with schizotypy. Following, we observed that individuals with higher positive schizotypal traits perceived self-generated touch with less sensory precision than individuals with lower positive schizotypal traits, without any effect on the precision of externally generated touches. This result indicates for the first time that high positive schizotypal traits are not accompanied by generic deficits in processing afferent somatosensory information but only self-generated somatosensory feedback and enforces the view that self-disorders lie at the core of the schizophrenia spectrum (Borda and Sass, 2015; Henriksen et al., 2021; Northoff et al., 2021; Postmes et al., 2014; Sass and Parnas, 2003). Critically, both in terms of attenuation and precision of self-generated touch, it was the positive dimension of schizotypy that drove the effects and not the negative or disorganized dimension. This parallels the negative association previously observed between somatosensory attenuation and the severity of hallucinations (Shergill et al., 2014), as well as the delusional ideation (Palmer et al., 2016; Teufel et al., 2010) and passivity experiences (Lemaitre et al., 2016) of nonclinical individuals.

These deficits in somatosensory attenuation and precision can fall within the scope of subtle neurological aberrations in sensorimotor performance (e.g. motor coordination, graphesthesia) (Buchanan and Heinrichs, 1989; Schröder et al., 1991), that are present with variable severity across the psychosis continuum (Chan et al., 2018; Gaha et al., 2015; Herold et al., 2021; Janssen et al., 2009; Mechri et al., 2010). These neurological soft signs have been repeatedly associated with the negative symptoms of schizophrenia and negative schizotypy in non-clinical individuals (Bombin et al., 2005; Chan et al., 2015; Cvetić et al., 2009; Hembram et al., 2014; Kaczorowski et al., 2009; Prikryl et al., 2012; Theleritis et al., 2012; Tosaro and Dazzan, 2005; Varambally et al., 2006; Whitty et al., 2006; Yazici et al., 2002), and less robustly with the positive and disorganized dimensions (Barkus et al., 2006; de Leede-Smith et al., 2017; Mechri et al., 2010; Ojagbemi et al., 2015). Instead, our data revealed a relationship of somatosensory attenuation and precision only with positive schizotypy, and not with the negative and the disorganized dimensions. Consequently, our findings suggest that somatosensory attenuation and precision constitute a special category of neurological soft signs that is specifically related to the self and the positive dimension of psychotic and psychotic-like symptoms.

Our results provide important insights for understanding the mechanism underlying the positive symptoms of schizophrenia. From a computational perspective, our effects can be explained by a deficit in the internal forward model that predicts the somatosensory consequences of the movement. Earlier studies have shown that somatosensory attenuation relies on spatiotemporal motor predictions (Bays et al., 2005; Bays and Wolpert, 2008; Kilteni et al., 2019) and not on postdictive processes (Bays et al., 2006; Kilteni and Ehrsson, 2020b), and it requires conditions where the received touch can be predicted by the motor command (Bays et al., 2006, 2005; Kilteni et al., 2021, 2020, 2018; Kilteni and Ehrsson, 2017b, 2017a; Shergill et al., 2003). In our study, the reduced attenuation indicates that with the same motor command, the brain of an individual with high positive schizotypy does not accurately predict the sensory consequences of the voluntary movement, and this leads to less attenuation of the self-generated somatosensory feedback compared to an individual with low positive schizotypy. Subsequently, the combination of this inaccurately predicted somatosensory information with the actual somatosensory feedback at the level of state estimation further leads to the decreased precision of self-generated touches. Within a Bayesian framework where prediction corresponds to prior expectations and sensory feedback to sensory evidence (Fletcher and Frith, 2009), our study indicates that high positive schizotypy is related to atypical prior expectations (generated by the internal forward model) and atypical combination of prior knowledge with sensory evidence (state estimation).

The cerebellum has been repeatedly implicated in predicting the sensory consequences of one’s own actions (Bays et al., 2006; Blakemore et al., 2001; Sarah J. Blakemore et al., 1999; Shadmehr et al., 2010, 2008; Tanaka et al., 2021; Wolpert DM, Miall RC et al., 1998), and we previously showed that somatosensory attenuation depends on the functional connectivity between the cerebellum and the primary and secondary somatosensory cortices (Kilteni and Ehrsson, 2020a): the stronger this corticocerebellar connectivity during self-generated touches compared to externally generated touches, the greater the somatosensory attenuation. Schizophrenia is also strongly associated with alterations in structural and functional cerebellar connectivity (Kim et al., 2021; Moberget and Ivry, 2019). Patients show impairments in cerebellar-mediated motor tasks (Bernard and Mittal, 2014), deficits in the integrity of the cerebellar white matter tracts (Kanaan et al., 2009; Kyriakopoulos and Frangou, 2009), abnormal resting-state cerebellar connectivity (Anteraper et al., 2021) with the cortex (Shinn et al., 2015), including frontoparietal (Repovs et al., 2011) and sensorimotor networks (Collin et al., 2011; D. J. Kim et al., 2020; Walther et al., 2017), decreased cerebellar connectivity with the primary motor cortex (Moussa-Tooks et al., 2019) and altered cerebellar activation (Bernard and Mittal, 2015) compared to healthy controls. Intriguingly, individuals at ultra-high-risk for psychosis have decreased resting-state cerebellocortical connectivity compared to controls (Bernard et al., 2014), while functional and structural cerebellocortical connectivity relates to their positive symptom progression (Bernard et al., 2017). Given these results, it was recently proposed to use cerebellocortical connectivity as a state-independent neural signature for psychosis prediction and characterization (Cao et al., 2018). Therefore, based on our findings, we speculate that positive schizotypy and consequently positive symptoms of schizophrenia are related to altered corticocerebellar connectivity in particular.

Future efforts should exploit the perception of self-generated somatosensation as a potential cognitive biomarker of psychosis. In contrast to other markers, including prepulse inhibition, mismatch negativity and P300 (Donati et al., 2020), which reflect deficits in processing externally generated information in schizophrenia, our results emphasize deficits in processing self-generated information. To date, there are no objective biological measures available to inform diagnostic or treatment decisions of schizophrenia (Martins-de-Souza, 2013; Waszkiewicz, 2020), hindering the early detection of disease onset in individuals who are at an increased risk for schizophrenia (Gottesman and Bertelsen, 1989; Tarbox and Pogue-Geile, 2011). To this end, attenuation and precision of self-generated somatosensation could reinforce the diagnostic procedure with an objective measure, which is meaningful since scale-based measures may be susceptible to self-report bias. Furthermore, given that the positive symptoms in the prodromal phase have high positive predictive power for the conversion of a high-risk state to schizophrenia (Klosterkötter, 2012; Meisenzahl et al., 2020), self-generated somatosensation could function as a neurocognitive marker that, when combined with other genetic, biochemical and neuroimaging markers (H. K. Kim et al., 2020; Kraguljac et al., 2021), forms a multilayered ‘signature’ for schizophrenia liability. So far, this perspective is still at a premature stage and the implementation in clinical settings is far from complete. Undoubtedly, appropriate clinical contextualization and validation through future longitudinal studies are necessary. Nonetheless, the present study suggests that deficits in processing self-generated tactile information can indicate increased liability for schizophrenia.

## Acknowledgments

E.A. was supported by the Åke Wibergs Foundation (Medical Research Grant M20-0038 granted to K.K.). X.J. and K.K. were supported by the Swedish Research Council (VR Starting Grant 2019-01909 granted to K.K.). Experimental costs were supported by the Swedish Research Council and the Strategic Research Area Neuroscience (StratNeuro Starting Grant granted to K.K.)

## Author contributions

K.K., X.J. and E.A. conceived and designed the experiment. X. J. and E. A. collected the data. K.K., E.A. and X.J. conducted the statistical analysis. E.A., K.K. and X.J. wrote the manuscript.

## Supplemental Figures

**Supplemental Figure S1.**
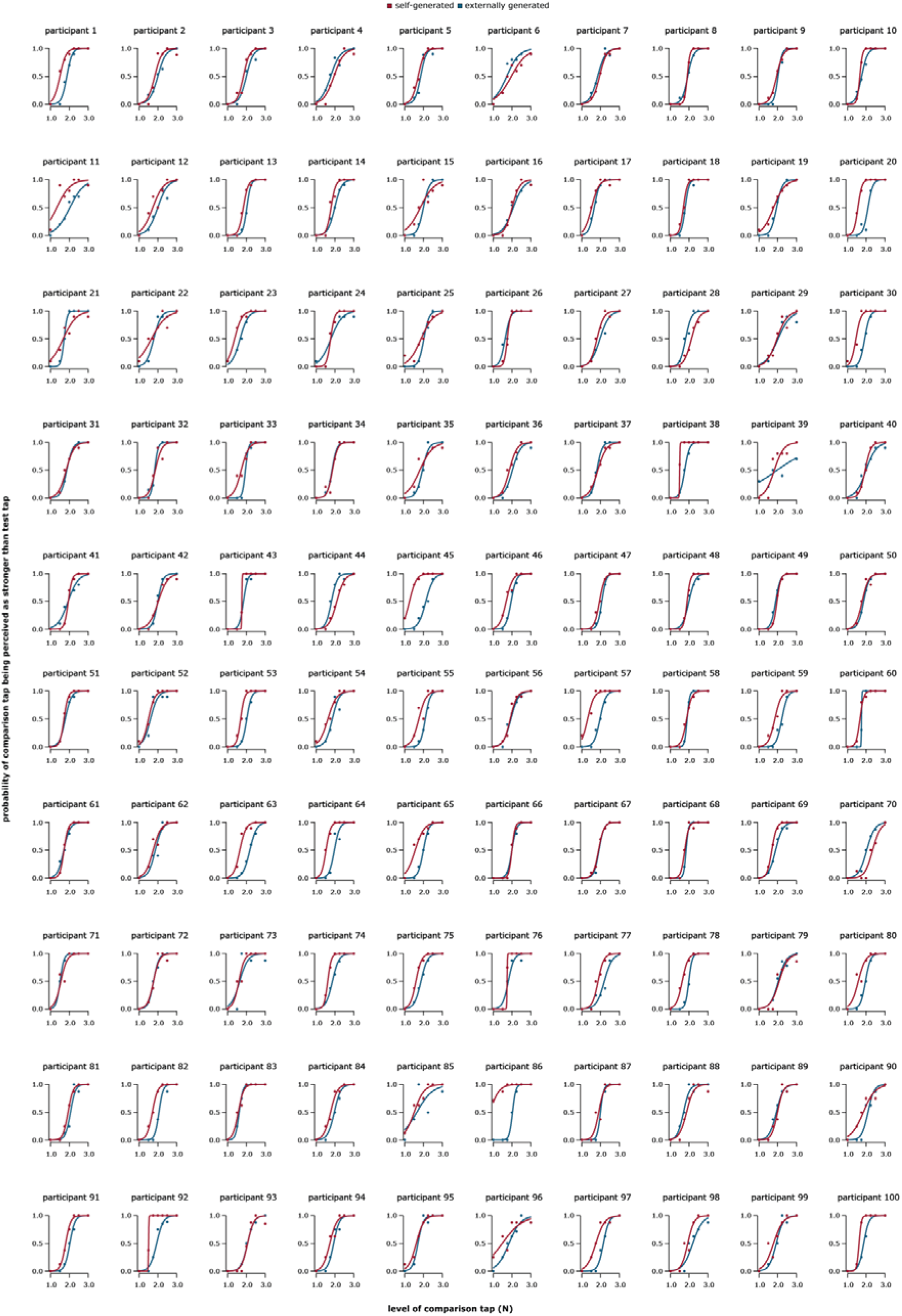
Fitted logistic models based on the participants’ responses under each condition.

**Supplemental Figure S2.**
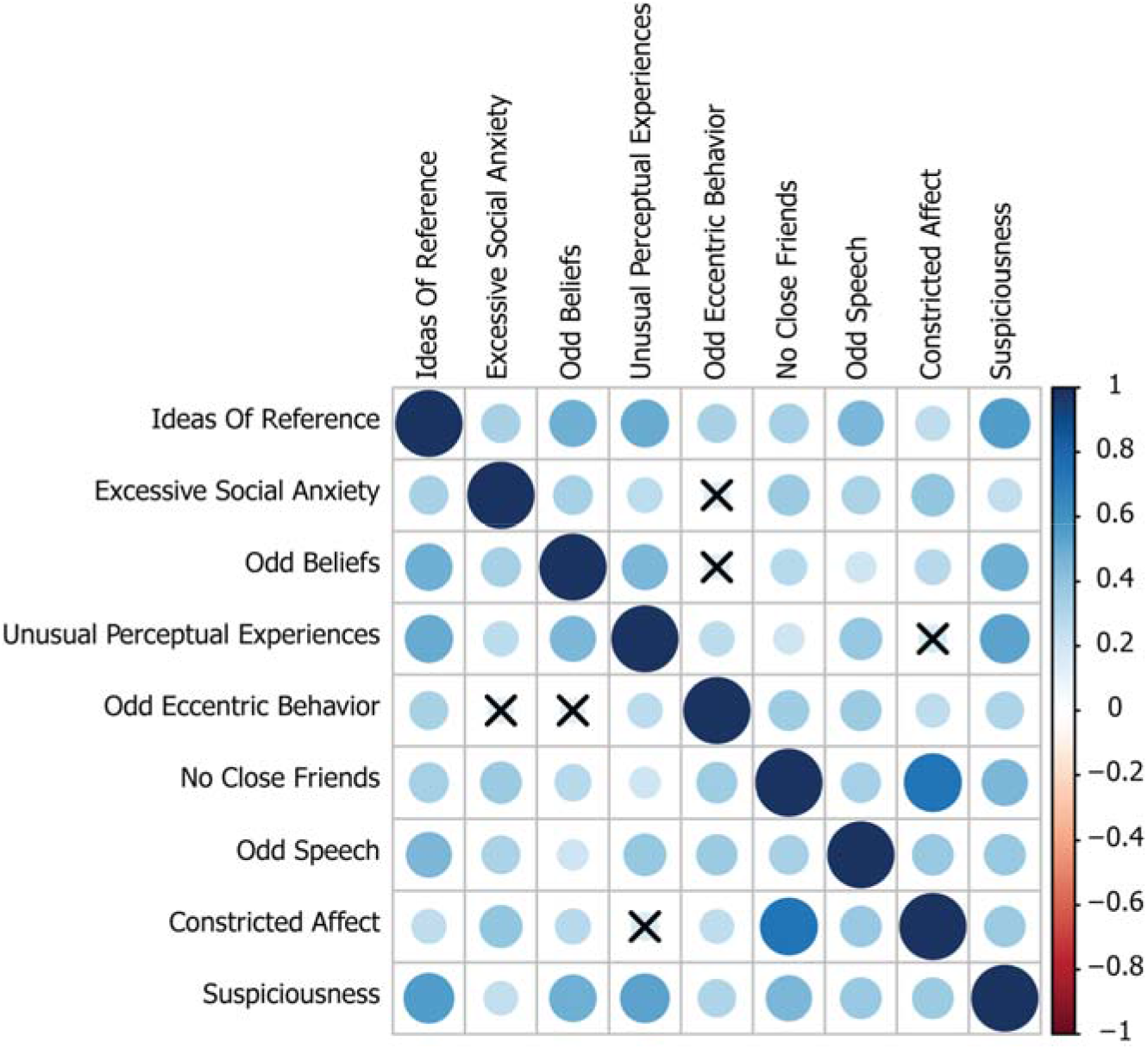
Correlations between the scores of all the SPQ subscales. The correlations have been calculated using the Spearman correlation coefficient and the *p*-values have been corrected for False Discovery Rate (FDR).

**Supplemental Figure S3.**
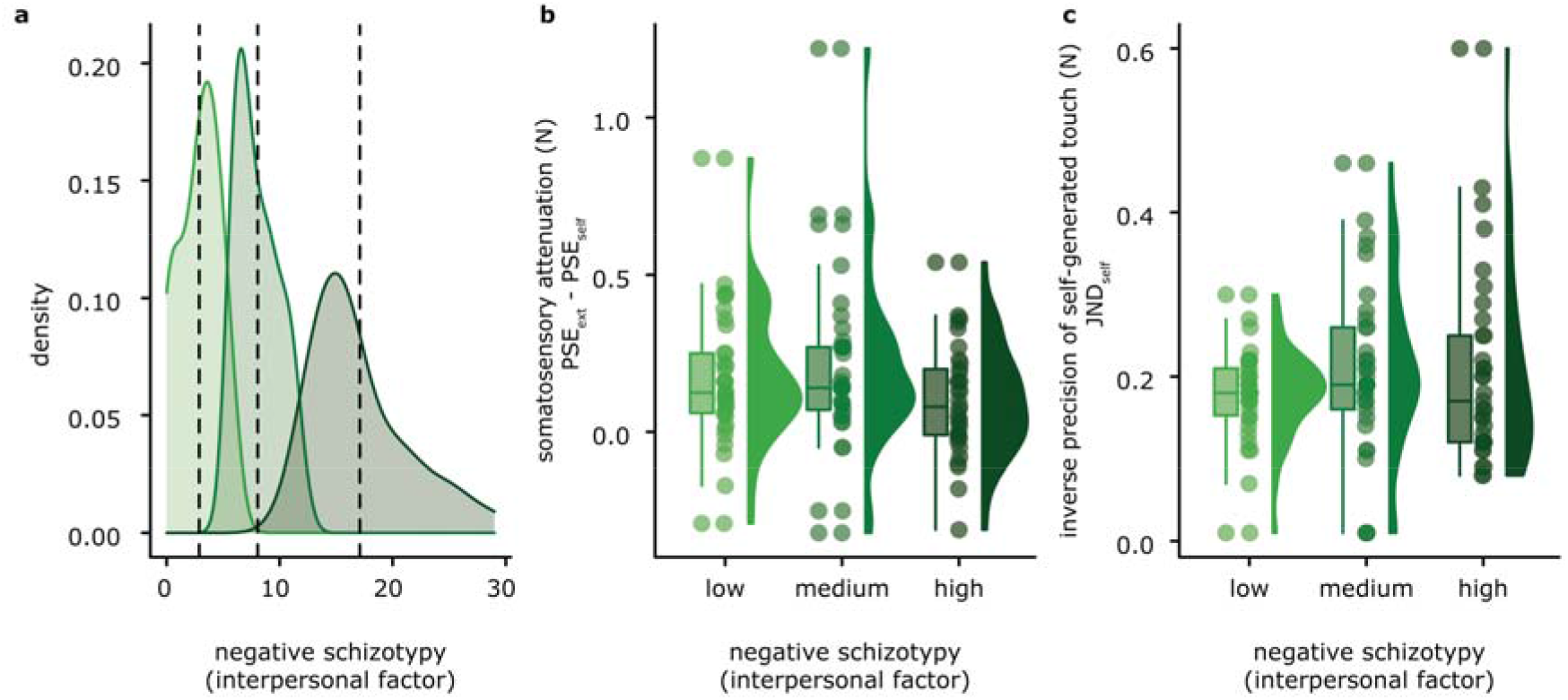
Somatosensory attenuation and precision in individuals with low, medium, and high negative schizotypal traits. **(a)** Density plots for the three schizotypy subgroups of our sample. Vertical dotted lines indicate the mean of each subgroup. **(b)** The boxplots show the median and interquartile ranges for somatosensory attenuation, the jittered points denote the raw data, and the violin plots display the full distribution of the data in each group. No significant differences in somatosensory attenuation were observed between the three groups. **(c)** The boxplots show the median and interquartile ranges for the JND in the *self-generated touch* condition, the jittered points denote the raw data, and the violin plots display the full distribution of the data in each group. No significant differences in somatosensory precision were observed between the three groups.

**Supplemental Figure S4.**
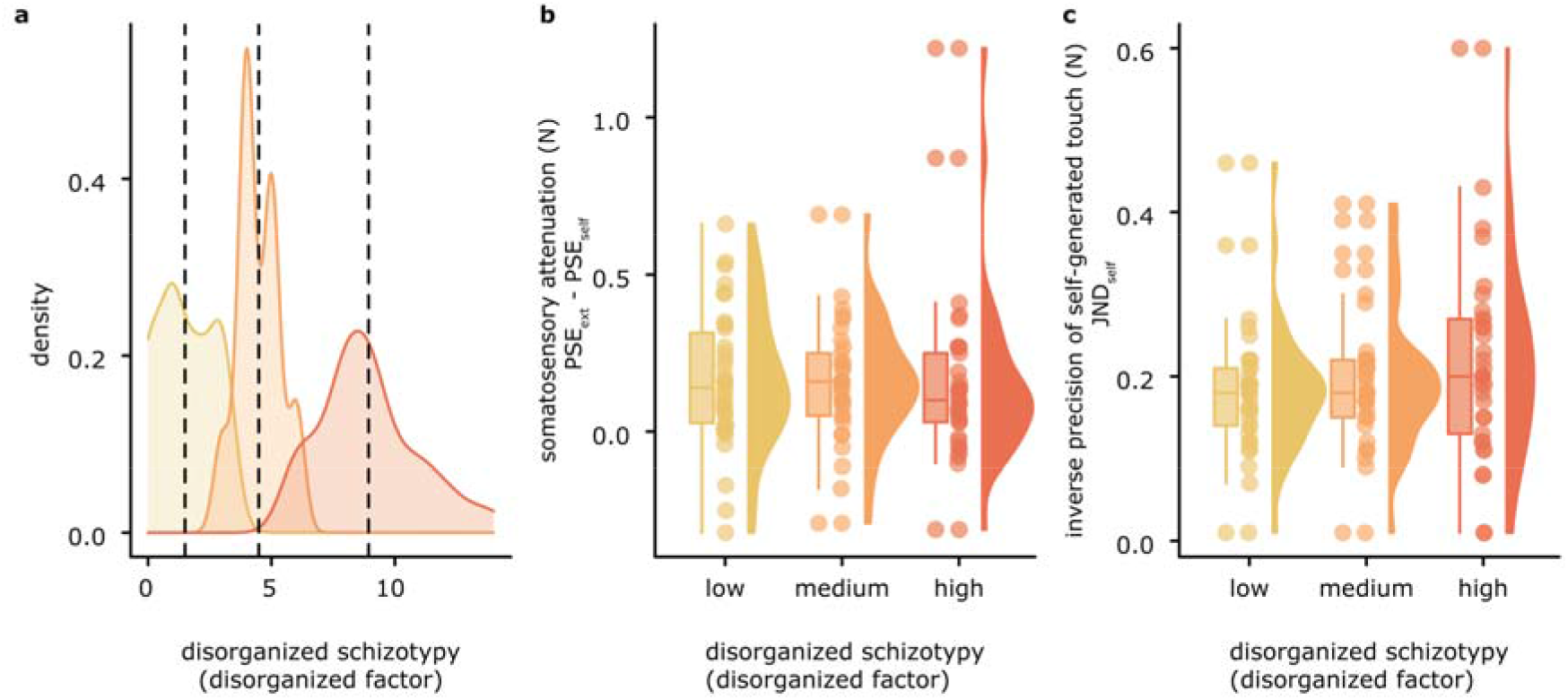
Somatosensory attenuation and precision in individuals with low, medium, and high disorganized schizotypal traits. **(a)** Density plots for the three schizotypy subgroups of our sample. Vertical dotted lines indicate the mean of each subgroup. **(b)** The boxplots show the median and interquartile ranges for somatosensory attenuation, the jittered points denote the raw data, and the violin plots display the full distribution of the data in each group. No significant differences in somatosensory attenuation were observed between the three groups. **(c)** The boxplots show the median and interquartile ranges for the JND in the *self-generated touch* condition, the jittered points denote the raw data, and the violin plots display the full distribution of the data in each group. No significant differences in somatosensory precision were observed between the three groups.

## Supplemental Tables

**Supplemental Table S1.** Descriptive statistics for the SPQ, its factors and its subscales.

| **SPQ Parameter** | **Mean** | **Standard Deviation** | **Variance** | **Range** |
| --- | --- | --- | --- | --- |
| *SPQ total* | 20.870 | 12.165 | 147.993 | 0-53 |
| **<u>Factors</u>** |  |  |  |  |
| *Cognitive-Perceptual* | 9.060 | 6.412 | 41.107 | 0-30 |
| *Interpersonal* | 9.280 | 6.566 | 43.113 | 0-29 |
| *Disorganized* | 4.940 | 3.381 | 11.431 | 0-14 |
| **<u>Subscales</u>** |  |  |  |  |
| *Ideas of Reference* | 3.130 | 2.489 | 6.195 | 0-9 |
| *Odd Beliefs or Magical Thinking* | 1.450 | 1.696 | 2.876 | 0-7 |
| *Unusual Perceptual Experiences* | 2.070 | 1.908 | 3.642 | 0-8 |
| *Odd or Eccentric Behaviour* | 1.630 | 1.868 | 3.488 | 0-6 |
| *Excessive Social Anxiety* | 2.870 | 2.460 | 6.054 | 0-8 |
| *No Close Friends* | 2.190 | 2.398 | 5.751 | 0-9 |
| *Odd Speech* | 3.310 | 2.214 | 4.903 | 0-9 |
| *Constricted Affect* | 1.810 | 1.756 | 3.085 | 0-7 |
| *Suspiciousness* | 2.410 | 2.000 | 4.002 | 0-8 |

## Supplemental Text

**Supplemental Text S1.**

Our sample had comparable levels of positive, negative and disorganized schizotypal traits, as can be seen in **Figure 2b-c**. A *Levene’s* test for Homogeneity of Variance revealed no significant difference between the variances of the positive and the negative schizotypy (cognitive-perceptual and interpersonal, *F*(1,198) = 0.115, *p* = 0.735). To compare the variances between the disorganized schizotypy scores and those of the cognitive perceptual or the interpersonal schizotypy, we first rescaled the disorganized scores (range: 0-16) to the same range as that of cognitive perceptual and interpersonal schizotypal scores (range: 0-33). We then performed the *Levene*’s test which revealed no significant difference in the variances of the cognitive-perceptual and the disorganized factors (*F*(1,198) = 0.691, *p* = 0.407), as well as between the interpersonal and disorganized factors (*F*(1,198) = 0.242, *p* = 0.624).

**Supplemental Text S2.**

To control whether the effects between somatosensory attenuation and precision are specific to the positive schizotypal traits, we repeated our categorical analysis for the negative and disorganized dimension of schizotypy.

***Negative schizotypy as a categorical variable***

We divided our sample into 3 subgroups: the low (*n_low_ = 34*), medium (*n_med_* = 33) and high (*n_high_* = 33) negative schizotypy groups (**Figure S2a**). There was no significant difference between the low and high schizotypy neither in terms of somatosensory attenuation (**Figure S2b**) (*n_low_ = 34, n_high_* = 33, *t*(63.5) = 1.484, *p* = 0.143, *CI^95^* = [−0.025, 0.167], *Cohen’s d* = 0.362, *BF_01_* = 1.58) nor in terms of somatosensory precision (*n_low_ = 34, n_high_* = 33, *W* = 561.5, *p* = 1, *CI^95^* = [−0.04, 0.04], *rrb* < 0.001, *BF_01_* = 3.680) (**Figure S2c**).

***Disorganized schizotypy as a categorical variable***

We divided our sample into 3 subgroups: the low (*n_low_ = 34*), medium (*n_med_* = 33) and high (*n_high_* = 33) disorganized schizotypy groups (**Figure S3a**). There was no significant difference between the low and high schizotypy neither in terms of somatosensory attenuation (**Figure S3b**) (*nlow = 34, n_high_* = 33, *W* = 628, *p* = 0.404, *CI^95^* = [−0.050, 0.130], *rrb* = 0.119, *BF_01_* = 3.368) nor in terms of somatosensory precision (*n_low_ = 34, n_high_* = 33, *W* = 460.5, *p* = 0.209, *CI^95^* = [−0.070, 0.010], *rrb* = −0.179, *BF_01_* = 2.389) (**Figure S3c**).

